# Helminth infection in a suburban ungulate population is driven more by age than landscape variables

**DOI:** 10.1101/2020.03.31.018531

**Authors:** J. Trevor Vannatta

**Affiliations:** University of Minnesota-Duluth, Swenson Science Building 207, 1035 Kirby Drive, Duluth, MN 55812 and Natural Resources Research Institute, University of Minnesota, 5013 Miller Trunk Highway, Duluth, Minnesota 55811-1442

**Keywords:** urban ecology, coinfection, infection risk, white-tailed deer, *Fascioloides magna*, *Taenia hydatigena*

## Abstract

Wildlife are increasingly common in suburban environments as towns and cities spread into surrounding rural areas. Many wildlife species have adapted to these new environments; however, we know comparatively little about how parasites respond urbanization of host habitats. Parasites are important members within ecological communities and alterations to transmission dynamics are known to alter host population dynamics. For complex life cycle parasites (parasites that use multiple different host species), suburban environments are thought to decrease transmission. Here, infection metrics of two parasites of white-tailed deer, giant liver flukes and thin-necked bladderworms, are examined to determine how successful these parasites are in a suburban environment. Additionally, land cover variables within suburban deer hunting areas are used to test if infection prevalence is associated with certain landscape level metrics. Results indicate that both parasites are common across the suburban landscape and are commonly found coinfecting the same hosts. Prevalence of neither parasite was strongly related to landscape variables within deer hunting areas, but fluke intensity was negatively correlated with the proportion of human development on the landscape. Overall, the scale of transmission events and host-parasite biology may explain why landscape metrics are weak predictors of infection risk in this system.

## INTRODUCTION

Many organisms adapt to urban and suburban landscapes (Crooks 2002; Magle et al. 2012; Brearley et al. 2013; Eakin et al. 2018), but as these organisms move into suburban habitats what happens to their parasites is less clear. Disease dynamics in these populations is often not considered (Bradley and Altizer 2007; Brearley et al. 2013; Murray et al. 2019) despite the clear ecological significance of host-parasite interactions (Tompkins et al. 2011; Buck and Ripple 2017; Vannatta and Minchella 2018). For complex life cycle parasites which use multiple different host species, it could be particularly difficult to predict how parasite populations will change in suburban ecosystems (Bradley and Altizer 2007; Brearley et al. 2013; Murray et al. 2019). Many complex life cycle parasites have cryptic intermediate hosts (e.g. amphibious vernal pool snails) which are difficult to find and sample. Additionally, complex life cycle parasites must have every host present in order to persist. A single missing host will lead to eradication of the parasite (Pickles et al. 2013; Murray et al. 2019). The multifaceted nature of these parasites makes predicting infection difficult especially as climate and other anthropogenic changes alter the landscape (Waller and Alverson 1997; Galatowitsch et al. 2009; Pickles et al. 2013; Dawe and Boutin 2016).

Landscape level cover type variables (as included in the National Land Cover Dataset) are often used to predict infection risk for complex life cycle parasites (Pickles et al. 2013; VanderWaal et al. 2015; Escobar et al. 2019), seemingly circumventing the need to identify the habitat of cryptic intermediate hosts. Alternatively, if cover types are associated with infection, researchers could use this information to find and adequately study these cryptic species. Despite the utility of landscape level habitat variables for predicting infection in other habitats, parasites in suburban ecosystems are rarely studied using these tools (Bradley and Altizer 2007; Brearley et al. 2013; Murray et al. 2019). Two complex life cycle parasites which can be used to explore these tools are the giant liver fluke (*Fascioloides magna*) and thin-necked bladderworm (*Taenia hydatigena*). Both parasites infect white-tailed deer (*Odocoileus virginianus*), a common resident of suburban landscapes. Both parasites have life cycle stages outside of a host (see below) which interact directly with the environment, and coinfection with both parasites is common. Liver flukes may contribute to mortality in cervids (Cheatum 1951; Murray et al. 2006; Shury et al. 2019), but often heavy fluke burdens have no apparent health effects (Pursglove et al. 1977; Lankester and Foreyt 2011; Wünschmann et al. 2015; Vannatta and Moen 2016). However, sublethal effects of liver fluke infections, such as decreased weight and number of antler tines in bucks during rut, and reduced twinning rates in does, are known (Mulvey and Aho 1993; Mulvey et al. 1994). Bladderworms are also largely innocuous (Blazek et al. 1985; Jones and Pybus 2001) except in extreme cases (Gregson 1937).

In the current study, metrics of giant liver fluke and thin-necked bladderworm infection are characterized within a suburban environment and compared to other studies from natural sites in the area. Additionally, habitat was classified within suburban deer hunting areas to determine whether utilization of landscape level tools can predict disease risk in a suburban landscape

## MATERIALS AND METHODS

### Study area

Duluth, Minnesota, USA is located on the western-most point of Lake Superior spanning an area of 176 km^2^. Duluth has a population of ∼86,000 people with a density of 489 people per km^2^ and 205 households per km^2^ placing the city within the classification of a suburb. Topography near the lake tends to be steep due to glacial uplift and becomes flatter farther from Lake Superior. The city contains many parks and hiking trails with vegetation predominantly classified as mixed deciduous and coniferous forest. Housing developments and residences are often immediately adjacent to city parks and deer hunting areas. Deer hunting areas consist of discrete management units positioned throughout the city. These polygons have been shaped to meet management goals and minimize logistical constraints (Fig. 1).

**Fig. 1.**
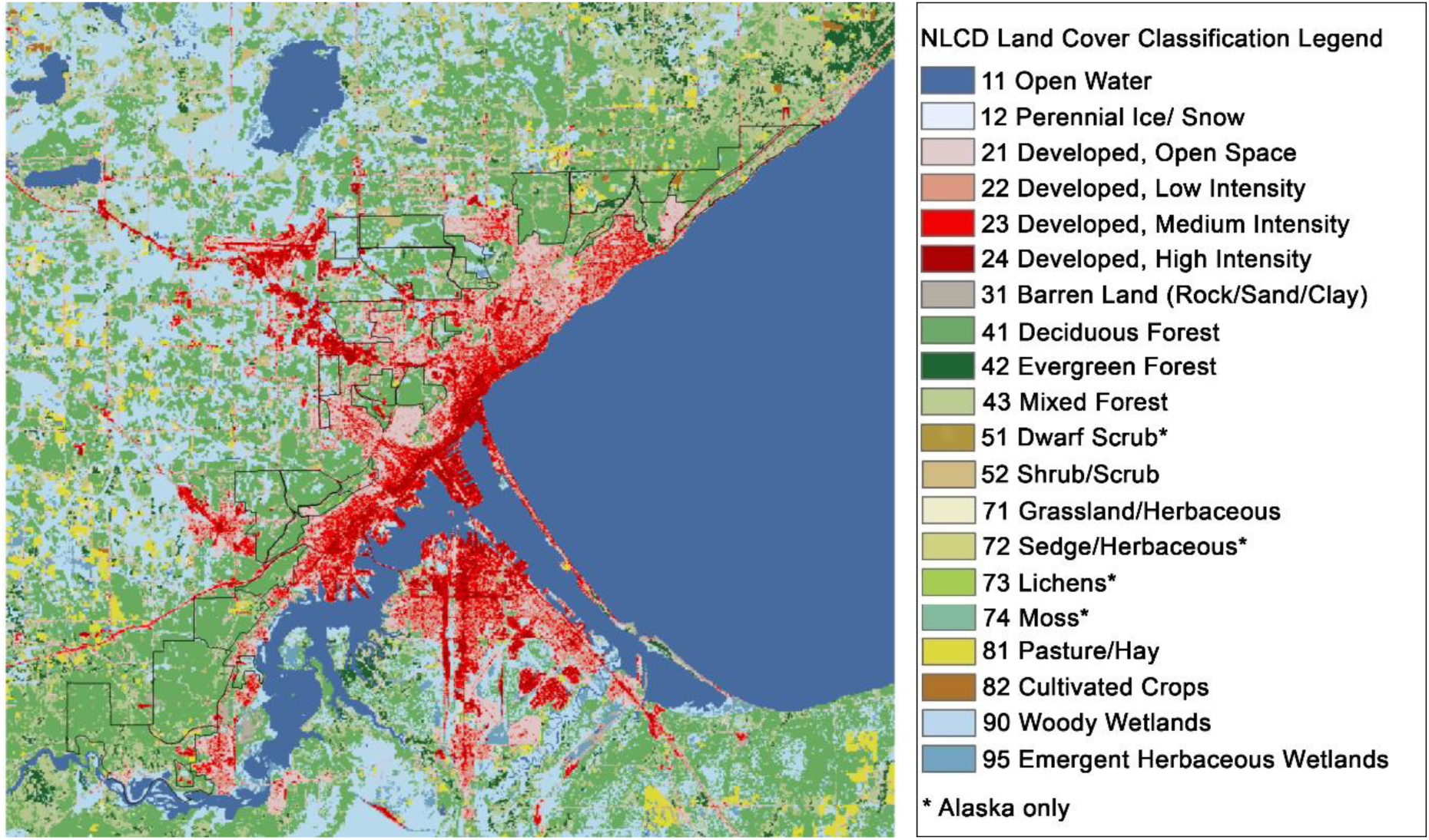
Deer hunting areas where samples were collected in the city of Duluth, Minnesota overlaid on the National Land Cover Dataset (2016). Deer hunting area polygons provided by the Arrowhead Bowhunters Alliance (www.bowhuntersalliance.org)

### Parasite life cycles

The giant liver fluke (*Fascioloides magna*) is a trematode parasite which infects white-tailed deer (*Odocoileus virginianus*), other cervids, and livestock in North America (Pybus 2001). Giant liver fluke eggs are released in deer fecal pellets. These eggs hatch in water and infect snails in the family Lymnaeidae. After asexual reproduction, parasites leave the snail and encyst on vegetation at the water’s edge where they are consumed by deer beginning the life cycle again (Pybus 2001). Many deer are also infected or coinfected with the thin-necked bladderworm (*Taenia hydatigena*), a cestode parasite (Addison et al. 1988). Thin-necked bladderworms infect various canids and felids in which the tapeworm matures and releases egg-filled proglottids in the predator’s scat. These proglottids carry eggs into the environment where they are ingested by foraging deer (Jones and Pybus 2001). Deer are then depredated or scavenged by predators. Canid and felid predators ingest the bladderworm cysts which develop to adult tapeworms in the predator’s intestine continuing the life cycle.

### Sample collection and processing

Livers (N = 274) were collected from deer killed by bowhunters during the months of September to December for five years (2014 – 2018) in the city of Duluth, Minnesota, USA (Fig. 1). Hunters are assigned to a single deer hunting area and must harvest an antlerless deer before taking an antlered animal. Hunters recorded the location of harvest and sex of each animal harvested. Livers were frozen until being thawed for processing. Thawed livers were weighed with mass used as a proxy for deer age (Parra et al. 2014) as bowhunters did not uniformly include age along with other sample information. Livers were then sectioned into 0.5 to 1.5 cm slices in order to count flukes, fluke capsules (Lankester and Foreyt 2011) and bladderworm cysts.

### Data analysis

I estimated prevalence, infection intensity (number of worms per animal), and parasite aggregation using the moment estimate *k* parameter as defined by Bliss and Fisher (1953). The *k* parameter is often used in macroparasite studies to describe the degree of parasite aggregation within hosts (Quinnell et al. 1995; Shaw et al. 1998). As the *k* parameter approaches infinity, the negative binomial distribution approximates a Poisson distribution (Bliss and Fisher 1953) with the number of parasites per host modelled as a Poisson process. As the value of *k* approaches 0, aggregation is more overdispersed with more parasites contained within fewer hosts (Shaw et al. 1998). Associations between liver fluke and bladderworm infections were assessed using Fisher’s exact test.

Due to the strong impact of liver mass on infection probability, generalized linear models were fit with liver mass as the sole predictor for both parasites. Goodness of fit was assessed for these models using the Hosmer and Lemeshow Goodness of Fit test for binary data and model performance was quantified by examining the area under the sensitivity/specificity curve. Liver mass was log transformed to meet model assumptions for this analysis.

For landscape analyses, hunters provided the deer hunting area from which each deer was harvested (N = 167 animals with location data across 29 deer hunting areas). Deer hunting area polygons and harvest information were retrieved from the Arrowhead Bowhunter’s Alliance (www.bowhuntersalliance.org), land cover data were taken from the National Land Cover Dataset (NLCD 2016), and elevation data was retrieved from National Elevation Data set (NED). Deer density within each hunting area was estimated by dividing the average number of animals collected within a hunting area by its area (deer harvested/ year/ km^2^). Since hunters provided only the deer hunting area and not specific coordinates of harvest, there is uncertainty in the exact habitat types experienced by deer within each hunting area. In order to better capture this variation, approximately 100 points were regularly distributed within each deer hunting area. These points were buffered to the size of an average suburban deer home range (∼0.63 km^2^; Bowman 2011). Point polygons were then used to calculate the proportion of cover types within each simulated deer home range. These values were then averaged within each deer hunting area and attributed to each animal collected within the deer hunting area. Cover types that were highly correlated or had less than 5% cover in deer hunting areas were either combined with like variables or omitted from further analysis. This resulted in five landscape variables: wet cover (open water, woody wetlands, and emergent wetlands), developed cover (developed open space, low, medium, and high intensity development), deciduous forest, evergreen and mixed forest, and low vegetation (shrub/scrub, herbaceous grasslands, cultivated crops, and hay/pasture). Barren land was omitted as this cover type had no comparable class with which it could be combined. This omission resulted in an average of 0.11% of deer hunting areas being omitted from the analysis. For elevation data, hunting area polygons were intersected with national elevation data and average polygon elevation and variance in elevation were calculated (Fig. S1). In Duluth, areas of high elevation are flatter and further from Lake Superior. Elevation variance was calculated to estimate vernal pool formation. Areas with low variance in elevation were expected to be flatter and more likely to contain vernal pools (as shown below using PCA), which support the intermediate hosts of giant liver flukes (Laursen et al. 1992; Laursen and Stromberg 1993). Due to remaining collinearity in predictor variables, deer hunting area attributes were subsequently input into a principal component analysis (PCA) to reduce dimensionality and prevent overfitting models. Following PCA, generalized linear mixed models were fit using liver mass and landscaped attributes as fixed variables (described above) and deer hunting area as a random factor. Additionally, negative binomial general linear mixed models were fit to predict infection intensity using the same variables. All analyses were run using R version 3.6.1 (R Core Team 2019). See supplementary code for packages used.

## RESULTS

Over five years, 274 deer livers were examined. The prevalence of liver flukes and bladderworms was 49% and 18%, respectively. Infection intensities for both parasites conformed to a negative binomial distribution, with flukes showing a higher degree of aggregation than bladderworms (Table 1 and Fig. 2). Coinfection with flukes and bladderworms was more common than expected by chance alone (12%; Fisher’s exact test p = 0.028; Table S1).

**Table 1.**
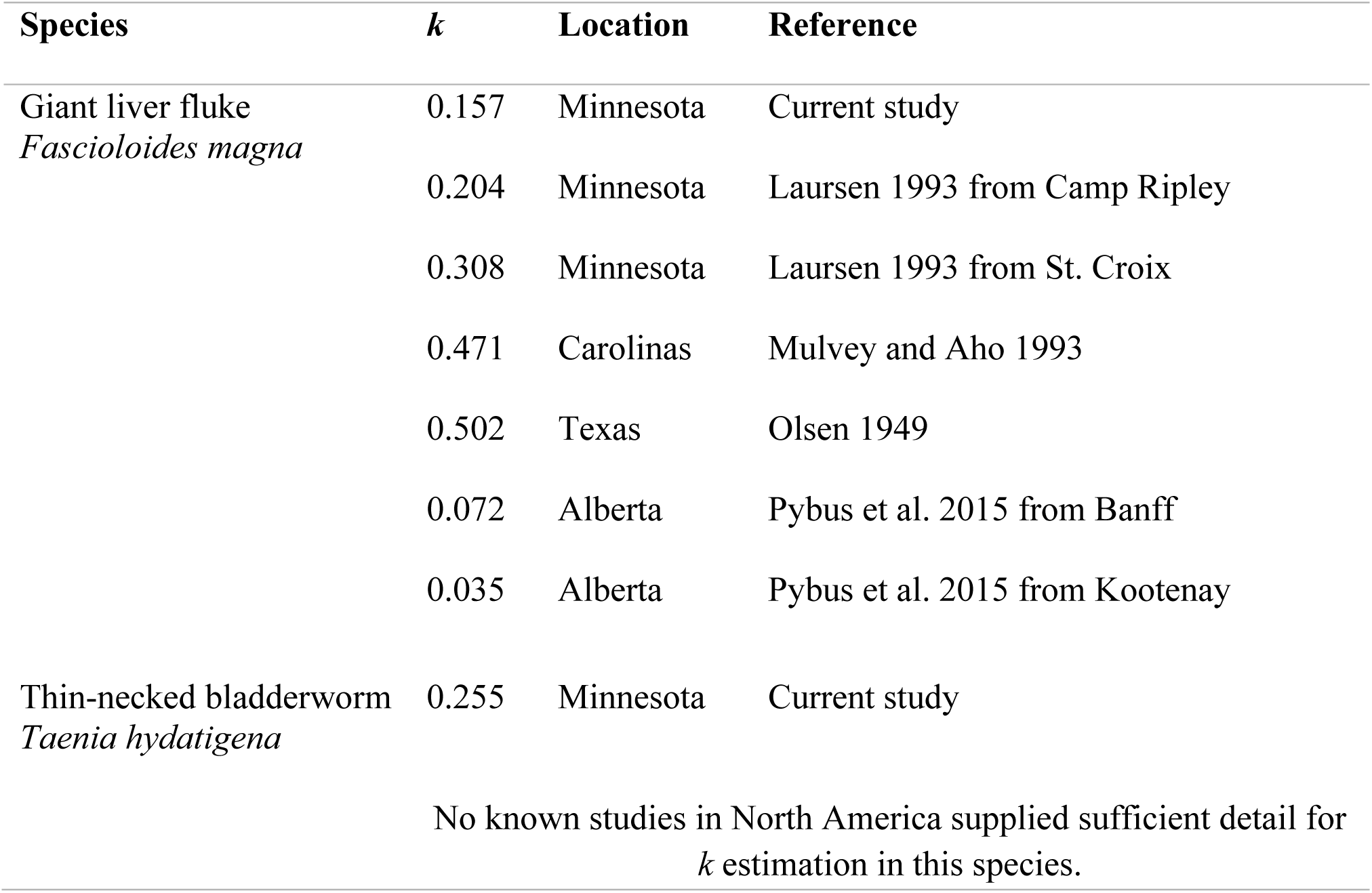
*k* estimates for giant liver flukes and thin-necked bladderworm in white-tailed deer from the current study and previous studies

**Fig. 2.**
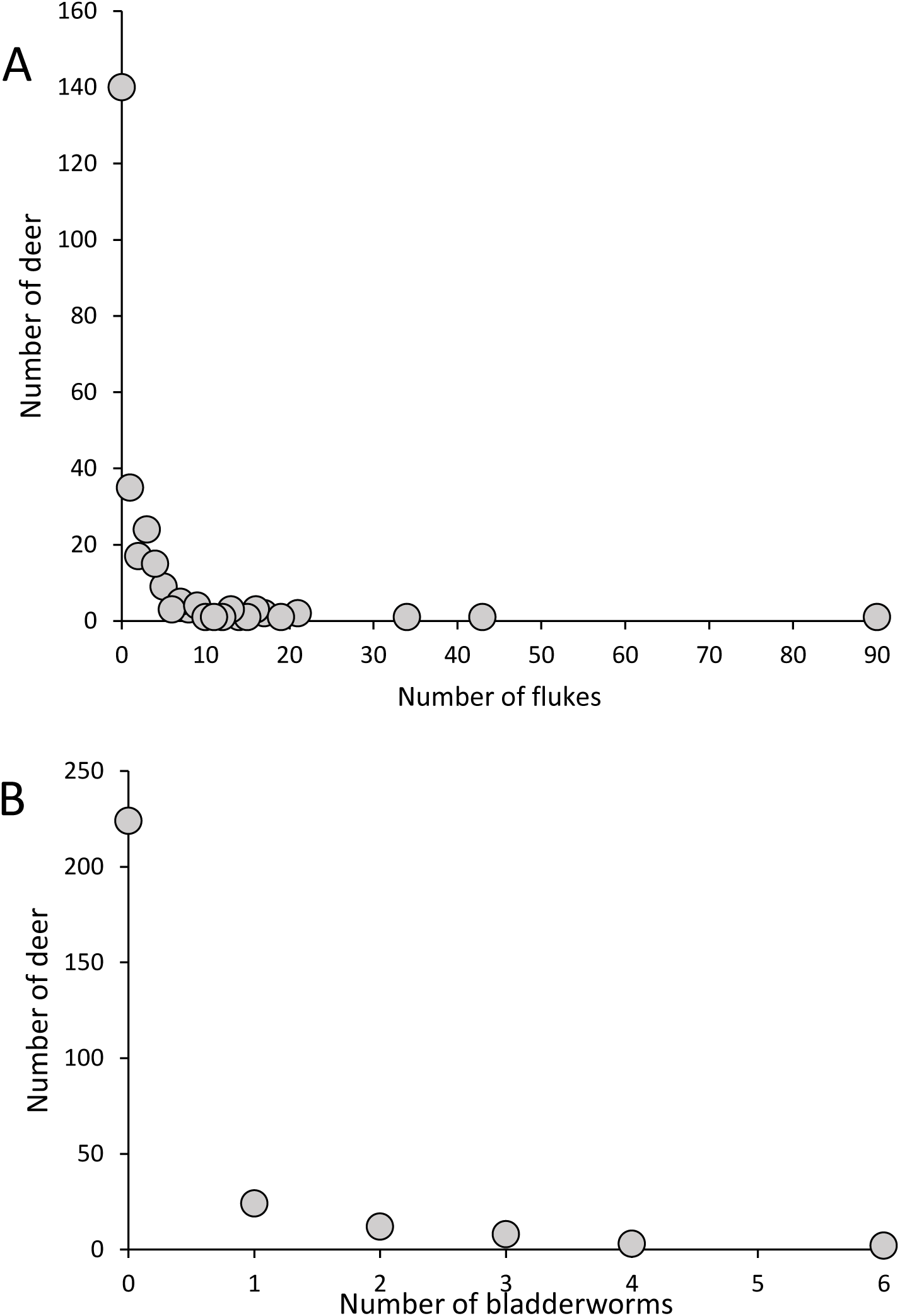
Frequency distributions for number of flukes (A) and number of bladderworms (B) within deer. Both parasites conform to a negative binomial distribution with *k* = 0.157 and 0.255 for flukes and bladderworms, respectively. This is equivalent to 53% of the fluke population being contained with 7% of the deer population, and 52% of the bladderworm population being contained within 5% of the deer population

Liver mass (a rough proxy for host age; Parra et al. 2014) was a significant predictor of infection status for both parasites (Fig. 3). Logistic regression models performed well (area under the curve (AUC) = 0.845 and 0.655 for fluke and bladderworm models, respectively) suggesting older animals with larger livers are more likely to be infected than younger animals. This trend also held for coinfections (Fig. S2).

**Fig. 3.**
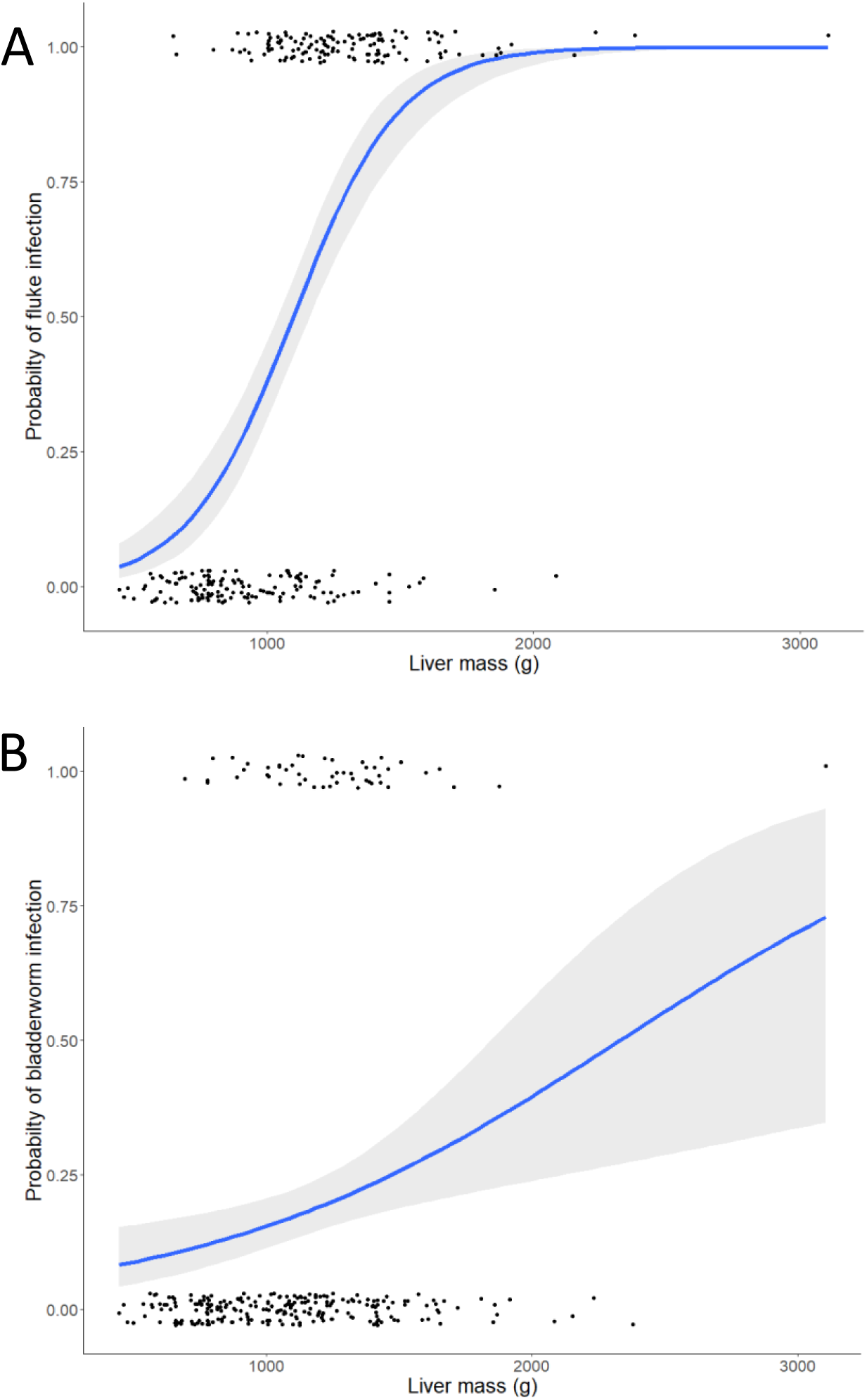
Logistic regression models for infection status as predicted by liver mass (age). Grey ribbon shows the 95% confidence interval of the curve. Untransformed liver mass is displayed for clarity. (A) The fluke prevalence curve performs well with an AUC of 0.845 (Log liver mass estimate = 5.684 ± 0.708, z = 8.024, p < 0.001).(B) The bladderworm curve, while still significant, performs less well with an AUC of 0.655 (Log liver mass estimate = 1.807 ± 0.543, z = 3.331, p = 0.001)

Multicollinearity was common within deer hunting area attributes (Fig. 4). PCA allowed for dimension reduction to two primary axes representing 52% of the data variance. PC1 was negatively correlated with suburban development and deer density (see table S2 for PC loadings). Additionally, PC1 was positively correlated with deciduous forest and hunting area size. Thus, PC1 was interpreted as the interaction between development, deer density, and habitat size. PC2 was positively correlated with elevation and wet cover types, and negatively correlated with evergreen/ mixed forest and the variance in elevation. Thus, PC2 represents the likelihood of inundation with areas of higher elevation (farther from Lake Superior) being flatter and wetter, and areas near the shore of Lake Superior being steeper with well drained soils supporting conifers and mixed forest.

**Fig. 4.**
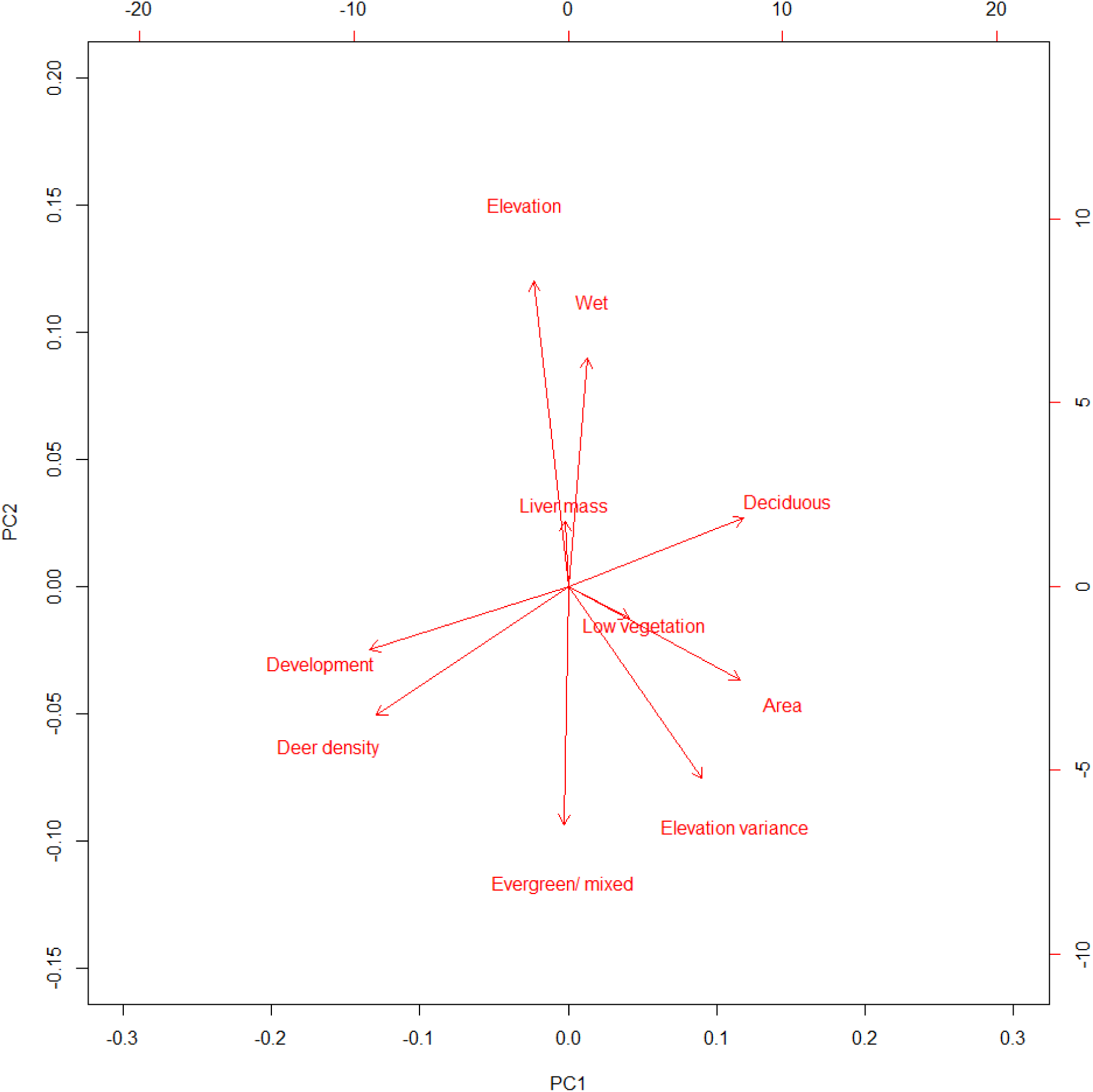
PCA showing multicollinearity among predictor variables. PC1 is interpreted as the interaction between development and deer density. PC2 represents the likelihood of inundation with areas of higher elevation (farther from Lake Superior) being flatter and wetter, and areas of lower elevation being steeper with well drained soils supporting conifers and mixed forest

Generalized linear mixed models containing liver mass (age) as a predictor variable for infection status consistently performed better than any model omitting this variable (Table 2, 3 and Table S3-S7). Additionally, no measure of habitat appeared more than once within the top models indicating these variables were not consistent predictors of infection. In models of infection intensity, bladderworm intensity is largely driven by liver mass (Table 3). In contrast, all three best performing fluke intensity models included liver mass and PC1 (Table 3).

**Table 2.**
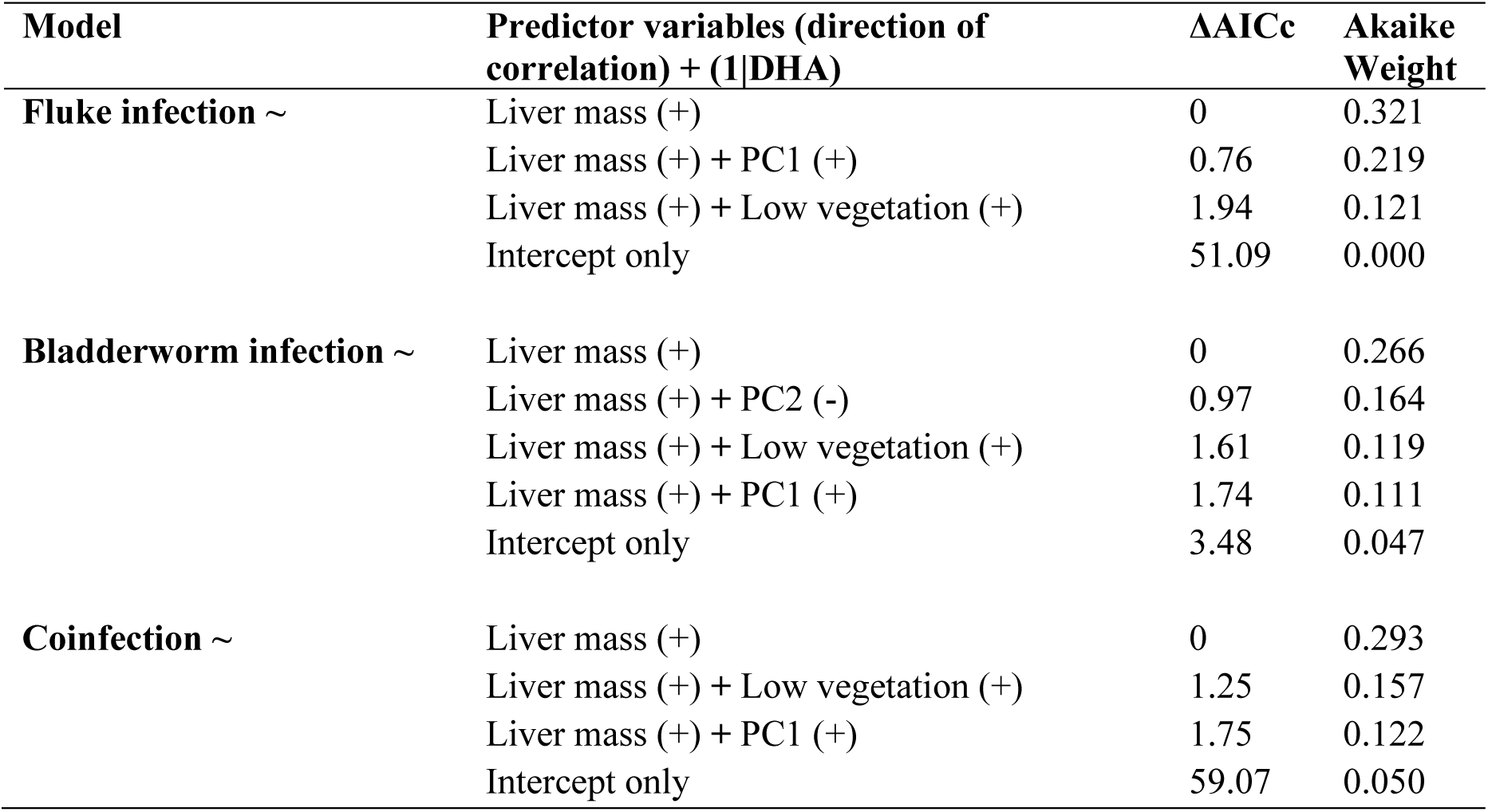
Generalized linear mixed models of infection probability for different infection types. Only intercept only models and models with ΔAIC less than 2 are displayed. All models include a random factor of deer hunting area + (1|DHA)

**Table 3.**
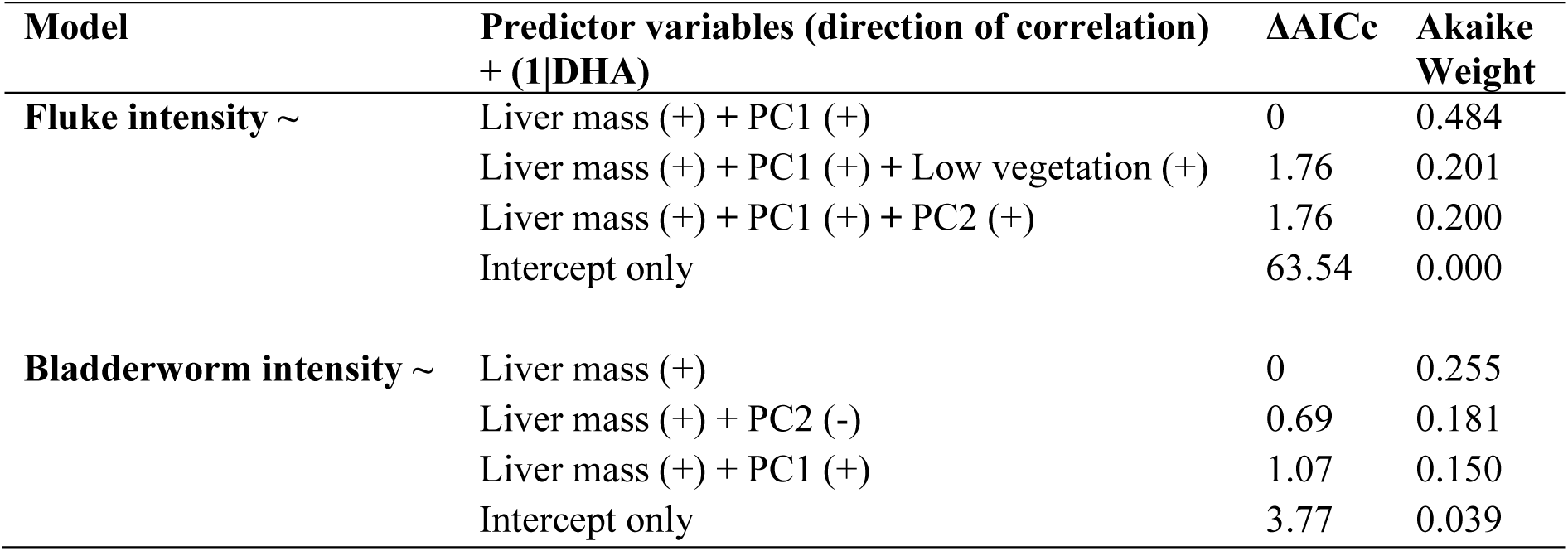
Negative binomial general linear mixed models of infection intensity for different infection types. Only intercept only models and models with ΔAIC less than 2 are displayed. All models include a random factor of deer hunting area + (1|DHA)

| Model | Predictor variables (direction of correlation) + (1\|DHA) | $\Delta AICc$ | Akaike Weight |
| --- | --- | --- | --- |
| **Fluke intensity ~** | Liver mass (+) + PC1 (+) | 0 | 0.484 |
|  | Liver mass (+) + PC1 (+) + Low vegetation (+) | 1.76 | 0.201 |
|  | Liver mass (+) + PC1 (+) + PC2 (+) | 1.76 | 0.200 |
|  | Intercept only | 63.54 | 0.000 |
| **Bladderworm intensity ~** | Liver mass (+) | 0 | 0.255 |
|  | Liver mass (+) + PC2 (-) | 0.69 | 0.181 |
|  | Liver mass (+) + PC1 (+) | 1.07 | 0.150 |
|  | Intercept only | 3.77 | 0.039 |

## DISCUSSION

In this study, I show that complex life cycle parasites fare well within suburban landscapes. Prevalence, intensity, aggregation, and coinfection of two helminth species, giant liver flukes (*Fascioloides magna*) and thin-necked bladderworm (*Taenia hydatigena*), were examined in suburban white-tailed deer. Measures of parasite prevalence and aggregation are largely similar to previous studies from natural areas in the region (see below) suggesting these parasites are adaptable to the suburban landscape of Duluth. Additionally, factors such as host age and landscape variables were used to predict complex life cycle helminth infections in suburban habitats. Models predicting infection probability strongly suggest that deer age is the primary driver of infection in this suburban deer population. Habitat attributes which have traditionally been associated with infection (wet cover types and deer density) did not strongly predict infection status in Duluth, although fluke intensity correlates with some landscape level features. This is likely a consequence of each parasite’s biology and the scale at which transmission occurs.

Many attributes of liver fluke parasitism reported here are consistent with other regional populations. The liver fluke prevalence in this study (49%) was similar to a study in natural areas of Ontario, Canada (52% Addison et al., 1988) and Voyageurs National Park, Minnesota (57% VanderWaal et al. 2015). However, the fluke prevalence seen currently in the Duluth area is ∼10-20% higher than the rest of northeastern Minnesota (Olsen and Fenstermacher 1943; Escobar et al. 2019). This contradicts previous work which suggested the Duluth area is suboptimal for fluke transmission compared to other regions in northern Minnesota, although this data was collected using fecal pellets in a population which likely experiences higher predation rates. (Escobar et al. 2019). Despite the suburban habitat of Duluth, the *k* parameter of giant liver fluke infection in the city is comparable to other studies from natural areas in Minnesota (see values from Laursen in Table 1). Combined, these values suggest not only that flukes can survive in the suburban environment, but that infection patterns largely align with natural areas. This similarity is likely related to the small scales at which transmission events occur (see below).

Bladderworm infection prevalence also aligns with other populations of white-tailed deer (Jones and Pybus 2001). Bladderworm prevalence in previous studies of the region (14% Olsen and Fenstermacher 1943) match the current study (18%). In contrast, bladderworms were present in 35% of deer in natural areas of Ontario, Canada (Addison et al. 1988). Although multiple factors likely contribute to the prevalence of these parasites, extirpation of carnivore hosts and range expansion of white-tailed deer in the region (Heffelfinger 2011; Dawe and Boutin 2016) are factors that could lead to fluctuations in parasite prevalence. In contrast to prevalence, no other studies provided sufficient detail to calculate *k* for other North American bladderworm populations (Table 1).

The correlation between parasite prevalence and liver mass is likely a result of host age since liver mass and deer age class (fawn, yearling, and adult) are correlated (Parra et al. 2014). Infection prevalence and intensity also increase with age for liver fluke (Cheatum 1951; Foreyt et al. 1977; Addison et al. 1988; Pybus 1990; Laursen 1993; Mulvey and Aho 1993; Qureshi et al. 1994) and bladderworm infections (Torgerson et al. 1995). This suggests parasitic infection and coinfection probability may increase with age for both parasites most likely because older animals have more time to become infected.

Although many studies have documented prevalence of liver flukes and bladderworms, I found none that reported the rate of coinfection. Coinfections have important ecological consequences and reporting parasite associations could be useful for understanding disease dynamics (Telfer et al. 2010). Many factors are thought to impact coinfection including environmental conditions, individual behavioral variation, and variation in immunity. My results suggested that host age (liver mass) was the only consistent predictor of coinfection (Table 2) likely for the same reasons mentioned above for individual infections.

Few studies have found landscape level correlates of liver fluke infection (Mulvey et al. 1994; Peterson et al. 2013; Pybus et al. 2015; VanderWaal et al. 2015; Escobar et al. 2019) and no studies of landscape influences on North American bladderworms were found during literature searches. Previous work on liver flukes has found swampy locations (Mulvey et al. 1994) and rooted-floating aquatic marshes to be significant predictors of fluke infections (VanderWaal et al. 2015). These conclusions likely relate to palatable aquatic plants and intermediate host habitat (Vannatta and Moen 2016). However, results have been mixed with some suggesting wetland habitat may be secondary to deer density for predicting infection risk (Peterson et al. 2013). The current study found no consistent landscape predictors of fluke prevalence (although liver mass was a significant predictor). However, fluke intensity was positively correlated with PC1 in all three best fitting models, suggesting higher intensity infections are found in larger deer hunting units, which contain more deciduous forest, less development, and lower deer densities. This result seems to contradict studies suggesting deer density as a positive driver of infection (Peterson et al. 2013), but this is likely related to multicollinearity. Deer thrive on habit edges created by development, which explains the positive correlations between these two variables. In contrast, where there is more development, deer hunting areas must be smaller and more fragmented explaining the negative correlation between development and area and deciduous forest. This result suggests that the scale at which individual transmission events are occurring is finer than traditional landscape level variables are able to distinguish. In suburban Duluth, the likely intermediate hosts of liver flukes are the *Fossaria* spp. snails, which are often vernal pool inhabitants (Vannatta and Moen 2016). Vernal pools can occur in a variety of cover types, such as deciduous forest, and on very small scales (Van Meter et al. 2008; Eakin et al. 2018). The significant associations found in previous studies may be due to broad, nonspecific cover type classifications (e.g. upland vs. swamp from Mulvey et al. 1994) or the presence of alternative intermediate hosts more restricted to permanent water bodies (possibly *Lymnaea elodes* in Voyageurs National Park, MN; VanderWaal et al. 2015). In contrast, the PC axis associated with deer density (as measured by harvest data) was not a significant predictor of fluke infection status in this study. This may be due to high densities of deer in all hunting areas driving prevalence to its regional maximum.

For bladderworm infections, the lack of landscape level predictors is most likely related to the high mobility of both deer and the definitive hosts, canid and felid predators, within their home ranges (Jones and Pybus 2001). A number of canid and felid species inhabit Duluth (coyotes, *Canis latrans*; red foxes, *Vulpes vulpes*; gray foxes, *Urocyon cinereoargentus*; domestic dogs, *C. famliaris*; wolves, *C. lupus*; and bobcat, *Lynx rufus*) and many of these species move through a suite of habitats on a regular basis (Crooks 2002; Ordenana et al. 2010; Mueller et al. 2018; Parsons et al. 2019). Additionally, canids/felids, deer, and even the bladderworm eggs disperse widely (Gemmell and Johnstone 1976; Gemmell et al. 1987; Torgerson et al. 1995) and are not restricted to specific habitat types. Thus, high dispersal within many cover types could explain why bladderworm infection was not associated with landscape attributes. Coinfection followed the same pattern as flukes and bladderworms did individually. This is not surprising as coinfection by necessity must relate to the attributes of single infection.

Dispersal of white-tailed deer presents a potential confounding variable in this study as deer may not reside permanently within the hunting area in which they are harvested. My data analysis techniques partly account for this by buffering regularly distributed points within each hunting area by suburban deer home range sizes. Additionally, suburban white-tailed deer often have home ranges much smaller than the average hunting area (0.6 km^2^ suburban deer home range (Bowman 2011) vs. 2.7 km^2^ average hunting area size). Lastly, dispersal rates of suburban white-tailed deer are low with one study only documenting a 10% dispersal rate (Clevinger 2017). Dispersal is also strongly male biased (Clevinger 2017) and hunter reported sex in our study suggests 65% of our livers were female which often have overlapping home ranges (Porter et al. 1991).

The current study monitored liver fluke and bladderworm infection and coinfection in a suburban deer population using landscape level tools to assess infection prevalence. Infection prevalence for both liver flukes and bladderworms align with previous studies. However, parasite aggregation (as measured by *k*) appears to be highly variable among populations, even within Minnesota. Landscape level variables were largely weak predictors of liver fluke and bladderworm infection status, likely related to the small scale of intermediate host habitat and the high mobility of hosts, respectively. This pattern held for coinfection as well. However, liver fluke infection intensity was negatively correlated with development and positively correlated with deciduous forest, likely related to intermediate host habitat. The additions of deer aging techniques, genetics/genomics, and GPS collaring of suburban deer would be valuable for a better understanding of these disease systems. The current study suggests that complex life cycle parasites can succeed within suburban landscapes, but that predicting these infections at such a fine scale will require refined techniques.

## ACKNOWLEDGMENTS

I am indebted to Brian Borkholder, the Duluth city bowhunters, and the Arrowhead Bowhunters’ Alliance for volunteering their deer livers for this study. I would like to acknowledge Dr. Ron Moen for supporting this work and allowing me independence to pursue these questions. Critical feedback on this manuscript was provided by Dr. Ron Moen, Dr. Dennis Minchella, and the Purdue graduate student and quantitative ecology communities. This work was funded by The University of Minnesota – Duluth, The Environmental and Natural Resources Trust Fund, The Clean Water, Land, and Legacy Amendment, and the Minnesota Zoo. The presentation of this work was also facilitated by an *Alces* Newcomer’s travel award. Lastly, I would like to thank Megan Gorder, Sara Carpenter, Olivia Lockyear, Bailey Pyle, Hannah Melchiorre, Ao Yu, Stephanie Gutierrez, and Heather Vannatta for assistance processing and storing deer livers.

## DECLARATIONS

### Funding

This work was funded by The University of Minnesota – Duluth, The Environmental and Natural Resources Trust Fund, The Clean Water, Land, and Legacy Amendment, and the Minnesota Zoo.

### Conflicts of interest/Competing interests

The author has no conflicts of interest to declare

### Code and data availability

All code and data used for analyses will be made publicly available on GitHub, URL: https://github.com/vanna006

## Notes

https://github.com/vanna006/Helminth-suburban-white-tailed-deer

## REFERENCES

Addison EM, Hoeve J, Joachim DG, McLachlin DJ (1988) *Fascioloides magna* (Trematoda) and *Taenia hydatigena* (Cestoda) from white-tailed deer. Can J Zool 66:1359–1364

Blazek K, Schramlova J, Hulinska D (1985) Pathology of the migration phase of *Taenia hydatigena* (Pallas, 1766) larvae. Folia Parasitol (Praha) 32:127–137

Bliss CI, Fisher RA (1953) Fitting the negative binomial distribution to biological data. Biometrics 9:176–200

Bowman JL (2011) Managing white-tailed deer: exurban, suburban, and urban environments. In: Hewitt, DG (ed) Biology and management of white-tailed deer. CRC Press, Florida, pp 599–622

Bradley CA, Altizer S (2007) Urbanization and the ecology of wildlife diseases. Trends Ecol Evol 22:95–102. https://doi.org/10.1016/j.tree.2006.11.001

Brearley G, Rhodes J, Bradley A, et al (2013) Wildlife disease prevalence in human-modified landscapes. Biol Rev 88:427–442. https://doi.org/10.1111/brv.12009

Buck JC, Ripple WJ (2017) Infectious agents trigger trophic cascades. Trends Ecol Evol 32:681–694. https://doi.org/10.1016/j.tree.2017.06.009

Cheatum EL (1951) Disease in relation to winter mortality of deer in New York. J Wildl Manage 15:216–220

Clevinger GB (2017) Effect of urbanization on the survival and movements of localized populations of white-tailed deer in southern Indiana. Dissertation, Ball State University

Crooks KR (2002) Relative sensitivities of mammalian carnivores to habitat fragmentation. Conserv Biol 16:488–502. https://doi.org/10.1046/j.1523-1739.2002.00386.x

Dawe KL, Boutin S (2016) Climate change is the primary driver of white-tailed deer (*Odocoileus virginianus*) range expansion at the northern extent of its range; land use is secondary. Ecol Evol 1–17. https://doi.org/10.1002/ece3.2316

Eakin CJ, Hunter ML, Calhoun AJK (2018) Bird and mammal use of vernal pools along an urban development gradient. Urban Ecosyst 21:1029–1041. https://doi.org/10.1007/s11252-018-0782-6

Escobar LE, Moen R, Craft ME, VanderWaal KL (2019) Mapping parasite transmission risk from white-tailed deer to a declining moose population. Eur J Wildl Res 65:1–11. https://doi.org/10.1007/s10344-019-1297-z

Foreyt WJ, Samuel WM, Todd AC (1977) *Fascioloides magna* in white-tailed deer (*Odocoileus virginianus*): observations on the pairing tendency. J Parasitol 63:1050–1052

Galatowitsch S, Frelich L, Phillips-Mao L (2009) Regional climate change adaptation strategies for biodiversity conservation in a midcontinental region of North America. Biol Conserv 142:2012–2022. https://doi.org/10.1016/j.biocon.2009.03.030

Gemmell MA, Johnstone PD (1976) Factors regulating tapeworm populations: dispersion of eggs of *Taenia hydatigena* on pasture. Ann Trop Med Parasitol 70:431–434

Gemmell MA, Lawson JR, Roberts MG (1987) Population dynamics in echinococcosis and cysticercosis: evaluation of the biological parameters of *Taenia hydatigena* and *T. ovis* and comparison with those of *Echinococcus granulosus*. Parasitology 94:161–180

Gregson JD (1937) Cysticercosis in deer. Parasitology 29:409

Heffelfinger JR (2011) Taxonomy, evolutionary history, and distribution. In: Hewitt, DG (ed) Biology and management of white-tailed deer. CRC Press, Florida, pp 3–42

Jones A, Pybus MJ (2001) Taeniasis and echinococcosis. In: Samuel, WM, Pybus, MJ, Kocan, AA (eds) Parasitic diseases of wild mammals, Iowa State University Press, Iowa, pp 150–192

Lankester MW, Foreyt WJ (2011) Moose experimentally infected with giant liver fluke (*Fascioloides magna*). Alces 47:9–15

Laursen JR (1993) Fascioloides magna in Minnesota: regional ecology of the fluke and serodiagnosis in cattle. Dissertation, University of Minnesota.

Laursen JR, Averbeck GA, Conboy GA, Stromberg BE (1992) Survey of pulmonate snails of central Minnesota I. Lymnaeidae. J Freshw Ecol 7:25–33

Laursen JR, Stromberg BE (1993) *Fascioloides magna* intermediate snail hosts: habitat preferences and infection parameters. J Parasitol 79:302–303

Magle SB, Hunt VM, Vernon M, Crooks KR (2012) Urban wildlife research: Past, present, and future. Biol Conserv 155:23–32. https://doi.org/10.1016/j.biocon.2012.06.018

Mueller MA, Drake D, Allen ML (2018) Coexistence of coyotes (*Canis latrans*) and red foxes (*Vulpes vulpes*) in an urban landscape. PLoS One 13:e0190971. https://doi.org/10.1371/journal.pone.0190971

Mulvey M, Aho JM (1993) Parasitism and mate competition: liver flukes in white-tailed deer. Oikos 66:187–192

Mulvey M, Aho JM, Rhodes E (1994) Parasitism and white-tailed deer: timing and components of female reproduction. Oikos 70:177–182

Murray DL, Cox EW, Ballard WB, et al (2006) Pathogens, nutritional deficiency, and climate influences on a declining moose population. Wildl Monogr 166:1–30

Murray MH, Sánchez CA, Becker DJ, et al (2019) City sicker? A meta-analysis of wildlife health and urbanization. Front Ecol Environ 17:575–583. https://doi.org/10.1002/fee.2126

Olsen OW (1949) White-tailed deer as a reservoir host of the large American liver fluke. Vet Med 44:26–30

Olsen OW, Fenstermacher R (1943) The helminths of North American deer with special reference to those of the white-tailed deer (*Odocoileus virginianus borealis*) in Minnesota. University of Minnesota Agricultural Experiment Station Technical Bulletin 159:1–20

Ordenana MA, Crooks KR, Boydston EE, et al (2010) Effects of urbanization on carnivore species distribution and richness. J Mammal 91:1322–1331. https://doi.org/10.1644/09-MAMM-A-312.1.Key

Parra CA, Duarte A, Luna RS, et al (2014) Body mass, age, and reproductive influences on liver mass of white-tailed deer (*Odocoileus virginianus*). Can J Zool 92:273–278

Parsons AW, Rota CT, Forrester T, et al (2019) Urbanization focuses carnivore activity in remaining natural habitats, increasing species interactions. J Appl Ecol 56:1894–1904. https://doi.org/10.1111/1365-2664.13385

Peterson WJ, Lankester MW, Kie JG, Bowyer RT (2013) Geospatial analysis of giant liver flukes among moose: effects of white-tailed deer. Acta Theor 58:359–365. https://doi.org/10.1007/s13364-013-0130-4

Pickles RSA, Thornton D, Feldman R, et al (2013) Predicting shifts in parasite distribution with climate change: a multitrophic level approach. Glob Chang Biol 19:2645–2654. https://doi.org/10.1111/gcb.12255

Porter WF, Mathews NE, Underwood HB, et al (1991) Social organization in deer: implications for localized management. Environ Manage 15:809–814. https://doi.org/10.1007/BF02394818

Pursglove SR, Prestwood AK, Ridgeway TR, Hayes FA (1977) *Fascioloides magna* infection in white-tailed deer of southeastern United States. J Am Vet Med Assoc 171:936–938

Pybus MJ (2001) Liver Flukes. In: Samuel, WM, Pybus, MJ, Kocan, AA (eds) Parasitic diseases of wild mammals, Iowa State University Press, Iowa, pp 121–149

Pybus MJ (1990) Survey of hepatic and pulmonary helminths of wild cervids in Alberta, Canada. J Wildl Dis 26:453–459

Pybus MJ, Butterworth EW, Woods JG (2015) An expanding population of the giant liver fluke (*Fascioloides magna*) in Elk (*Cervus canadensis*) and Other Ungulates in Canada. J Wildl Dis 51:431–445. https://doi.org/10.7589/2014-09-235

Quinnell RJ, Grafen A, Woolhouse MEJ (1995) Changes in parasite aggregation with age: a discrete infection model. Parasitology 111:635–644

Qureshi T, Lynn D, Davis DS, Craig TM (1994) Use of bait containing triclabendazole to treat *Fascioloides magna* infections in free ranging white-tailed deer. J Wildl Dis 30:346–350

R Core Team (2019) R: A language and environment for statistical computing. R Foundation for Statistical Computing, Vienna, Austria URL: https://www.R-project.org/

Shaw DJ, Grenfell BT, Dobson AP (1998) Patterns of macroparasite aggregation in wildlife host populations. Parasitology 117:597–608. https://doi.org/10.1017/S0031182098003448

Shury TK, Pybus MJ, Nation N, et al (2019) *Fascioloides magna* in Moose (*Alces alces*) from Elk Island National Park, Alberta. Vet Pathol 56:476–485. https://doi.org/10.1177/0300985818823776

Telfer S, Lambin X, Birtles R, et al (2010) Species interactions in a parasite community drive infection risk in a wildlife population. Science 330:243–246

Tompkins DM, Dunn AM, Smith MJ, Telfer S (2011) Wildlife diseases: From individuals to ecosystems. J Anim Ecol 80:19–38. https://doi.org/10.1111/j.1365-2656.2010.01742.x

Torgerson PR, Pilkington J, Gulland FMD, Gemmell MA (1995) Further evidence for the long distance dispersal of taeniid eggs. Int J Parasitol 25:265–267

Van Meter R, Bailey LL, Campbell Grant EH (2008) Methods for estimating the amount of vernal pool habitat in the northeastern United States. Wetlands 28:585–593

VanderWaal KL, Windels SK, Olson BT, et al (2015) Landscape influence on spatial patterns of meningeal worm and liver fluke infection in white-tailed deer. Parasitology 142:706–718. https://doi.org/10.1017/S0031182014001802

Vannatta JT, Minchella DJ (2018) Parasites and their impact on ecosystem nutrient cycling. Trends Parasitol 34:452–455. https://doi.org/10.1016/j.pt.2018.02.007

Vannatta JT, Moen R (2016) Giant liver fluke and moose: just a fluke? Alces 52:117–139

Waller DM, Alverson S (1997) The white-tailed deer: a keystone herbivore. Wildl Soc Bull 25:217–226

Wünschmann A, Armien AG, Butler E, et al (2015) Necropsy Findings in 62 Opportunistically Collected Free-Ranging Moose (*Alces Alces*) From Minnesota, USA (2003–13). J Wildl Dis 51:157–165. https://doi.org/10.7589/2014-02-037

