## Supplemental information for "Helminth infection in a suburban ungulate population is driven more by age than landscape variables"

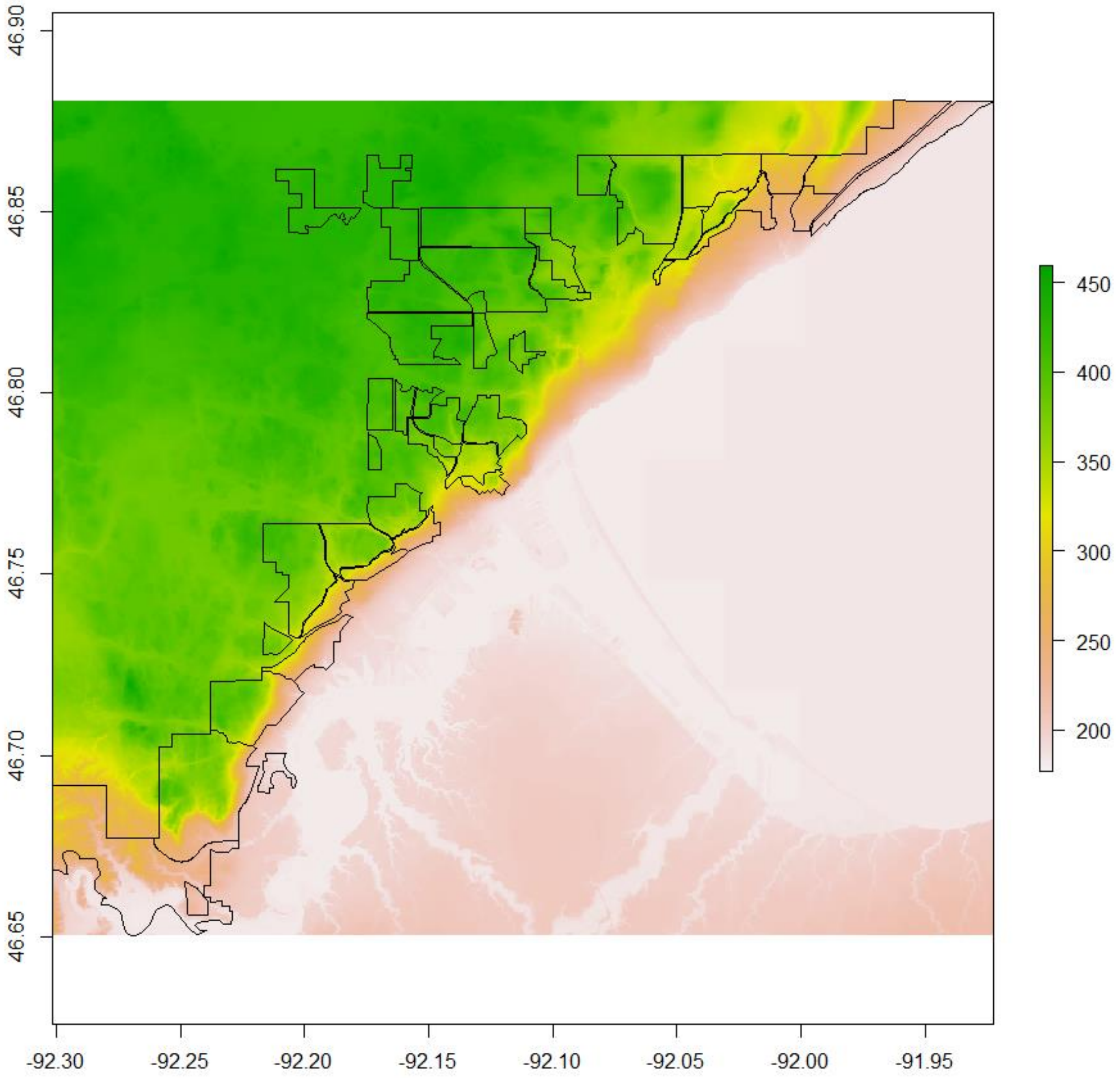


**Fig. S1** Deer hunting areas in Duluth, Minnesota over laid on the National Elevation Dataset. Elevation is in meters. Areas near the lake have lower elevations. Many hunting areas have high variation in elevation.

**Table S1** Contingency table of fluke and bladderworm infection. Coinfection was more common than expected by chance alone (Fisher's exact test $p = 0.028$)

| | *Bladderworm +* | *Bladderworm -* |
|---|---|---|
| *Fluke +* | 32 | 103 |
| *Fluke -* | 18 | 121 |

**Table S2** Results from principal component analysis with variable loadings

| **Landscape variable** | **PC1** | **PC2** |
|---|---|---|
| Proportion of variance explained | 0.3256 | 0.1928 |
| | Loadings | |
| Liver mass | -0.0090 | 0.1241 |
| Area | 0.4291 | -0.1775 |
| Deer density | -0.4806 | -0.2440 |
| Elevation | -0.0875 | 0.5795 |
| Elevation variance | 0.3347 | -0.3634 |
| Wet cover | 0.0465 | 0.4332 |
| Development | -0.4968 | -0.1186 |
| Deciduous forest | 0.4390 | 0.1302 |
| Evergreen and mixed forest | -0.1815 | 0.4781 |
| Low vegetation | -0.1769 | 0.2042 |

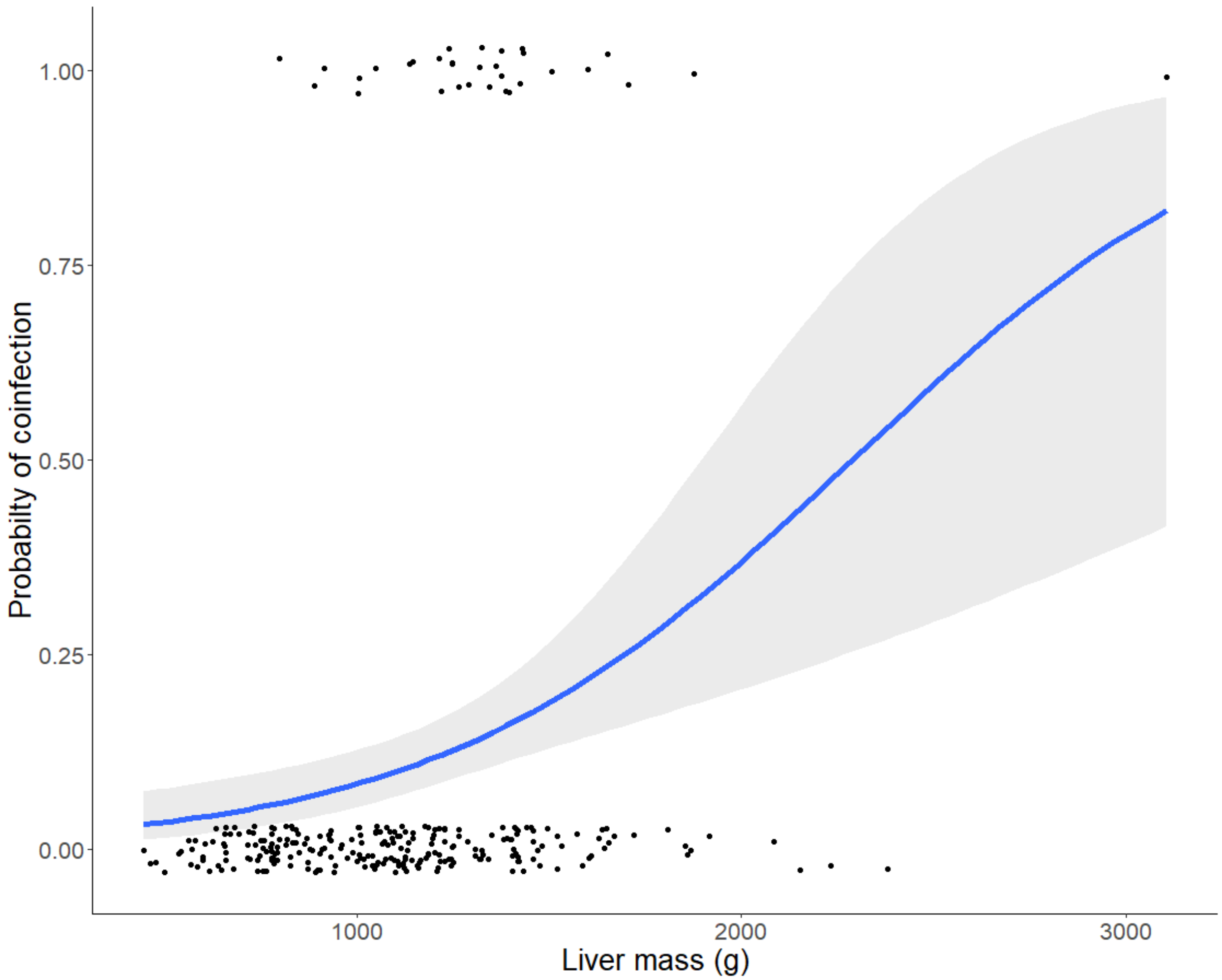


Fig. S2 Logistic regression model for coinfection status as predicted by liver mass (age). Black circles show actual data points with 0 being un of singly infected and 1 being coinfected. The coinfection curve performs well with an Area under the curve of 0.729 (Log liver mass coefficient = 2.7089 ± 0.6974, $z = 3.665$, $p = 0.001$).

**Table S3** Complete binomial model set for predicting liver fluke infection status. Blank values are omitted from the model. Model coefficients are presented under the variable. All models predict fluke infection status ~ present variables

| (Intercept) | Low vegetation | Liver mass | PC1 | PC2 | df | logLik | AICc | delta | weight |
|---|---|---|---|---|---|---|---|---|---|
| 0.12 | | 1.54 | | | 3 | -89.15 | 184.44 | 0 | 0.3208 |
| 0.11 | | 1.54 | 0.22 | | 4 | -88.48 | 185.20 | 0.76 | 0.2194 |
| 0.12 | 0.07 | 1.52 | | | 4 | -89.06 | 186.38 | 1.94 | 0.1217 |
| 0.12 | | 1.53 | | 0.04 | 4 | -89.13 | 186.50 | 2.06 | 0.1145 |
| 0.11 | | 1.54 | 0.22 | 0.04 | 5 | -88.45 | 187.27 | 2.83 | 0.0777 |
| 0.11 | 0.02 | 1.54 | 0.22 | | 5 | -88.47 | 187.31 | 2.88 | 0.0761 |
| 0.12 | 0.08 | 1.52 | | 0.04 | 5 | -89.04 | 188.45 | 4.01 | 0.0432 |
| 0.11 | 0.02 | 1.53 | 0.22 | 0.04 | 6 | -88.44 | 189.41 | 4.97 | 0.0267 |
| 0.04 | | | | | 2 | -115.73 | 235.53 | 51.09 | 2.58E-12 |
| 0.04 | | | | 0.19 | 3 | -115.02 | 236.19 | 51.75 | 1.86E-12 |
| 0.04 | | | 0.16 | | 3 | -115.20 | 236.54 | 52.10 | 1.56E-12 |
| 0.04 | 0.16 | | | | 3 | -115.21 | 236.56 | 52.12 | 1.54E-12 |
| 0.04 | 0.18 | | | 0.20 | 4 | -114.38 | 237.01 | 52.57 | 1.23E-12 |
| 0.04 | | | 0.16 | 0.19 | 4 | -114.48 | 237.21 | 52.77 | 1.11E-12 |
| 0.04 | 0.13 | | 0.13 | | 4 | -114.90 | 238.05 | 53.62 | 7.30E-13 |
| 0.04 | 0.15 | | 0.12 | 0.20 | 5 | -114.10 | 238.57 | 54.13 | 5.65E-13 |

**Table S4** Complete binomial model set for predicting bladderworm infection status. Blank values are omitted from the model. Model coefficients are presented under the variable. All models predict bladderworm infection status ~ present variables

| (Intercept) | Low vegetation | Liver mass | PC1 | PC2 | df | logLik | AICc | delta | weight |
|---|---|---|---|---|---|---|---|---|---|
| -1.62 | | 0.47 | | | 3 | -75.77 | 157.69 | 0.00 | 0.2660 |
| -1.63 | | 0.52 | | -0.25 | 4 | -75.21 | 158.66 | 0.97 | 0.1635 |
| -1.63 | 0.14 | 0.47 | | | 4 | -75.52 | 159.29 | 1.61 | 0.1191 |
| -1.61 | | 0.47 | 0.15 | | 4 | -75.59 | 159.43 | 1.74 | 0.1113 |
| -1.64 | 0.12 | 0.51 | | -0.24 | 5 | -75.02 | 160.40 | 2.72 | 0.0683 |
| -1.62 | | 0.52 | 0.14 | -0.23 | 5 | -75.04 | 160.46 | 2.77 | 0.0665 |
| -1.54 | | | | | 2 | -78.54 | 161.16 | 3.48 | 0.0468 |
| -1.62 | 0.11 | 0.47 | 0.11 | | 5 | -75.44 | 161.24 | 3.56 | 0.0449 |
| -1.62 | 0.10 | 0.51 | 0.11 | -0.23 | 6 | -74.93 | 162.38 | 4.70 | 0.0254 |
| -1.55 | 0.15 | | | | 3 | -78.25 | 162.65 | 4.97 | 0.0222 |
| -1.54 | | | | -0.13 | 3 | -78.38 | 162.91 | 5.22 | 0.0195 |
| -1.53 | | | 0.11 | | 3 | -78.43 | 163.01 | 5.32 | 0.0186 |
| -1.55 | 0.14 | | | -0.12 | 4 | -78.12 | 164.48 | 6.79 | 0.0089 |
| -1.54 | 0.13 | | 0.07 | | 4 | -78.21 | 164.67 | 6.99 | 0.0081 |
| -1.53 | | | 0.11 | -0.12 | 4 | -78.27 | 164.78 | 7.09 | 0.0077 |
| -1.54 | 0.12 | | 0.07 | -0.11 | 5 | -78.07 | 166.52 | 8.83 | 0.0032 |

**Table S5** Complete binomial model set for predicting coinfection status. Blank values are omitted from the model. Model coefficients are presented under the variable. All models predict coinfection status ~ present variables

| (Intercept) | Low vegetation | Liver mass | PC1 | PC2 | df | logLik | AICc | delta | weight |
|---|---|---|---|---|---|---|---|---|---|
| -2.26 | | 0.56 | | | 3 | -56.27 | 118.69 | 0.00 | 0.2927 |
| -2.25 | 0.21 | 0.56 | | | 4 | -55.85 | 119.94 | 1.25 | 0.1568 |
| -2.20 | | 0.56 | 0.20 | | 4 | -56.10 | 120.45 | 1.75 | 0.1218 |
| -2.25 | | 0.57 | | -0.06 | 4 | -56.25 | 120.75 | 2.06 | 0.1045 |
| -2.20 | 0.18 | 0.56 | 0.15 | | 5 | -55.76 | 121.89 | 3.20 | 0.0591 |
| -2.25 | 0.21 | 0.56 | | -0.04 | 5 | -55.84 | 122.04 | 3.35 | 0.0548 |
| -2.13 | | | | | 2 | -59.07 | 122.22 | 3.53 | 0.0502 |
| -2.19 | | 0.57 | 0.20 | -0.06 | 5 | -56.07 | 122.52 | 3.82 | 0.0433 |
| -2.11 | 0.21 | | | | 3 | -58.62 | 123.39 | 4.70 | 0.0279 |
| -2.20 | 0.17 | 0.57 | 0.15 | -0.05 | 6 | -55.74 | 124.01 | 5.31 | 0.0205 |
| -2.10 | | | 0.11 | | 3 | -59.00 | 124.15 | 5.46 | 0.0191 |
| -2.14 | | | | 0.09 | 3 | -59.03 | 124.20 | 5.51 | 0.0186 |
| -2.13 | 0.22 | | | 0.09 | 4 | -58.57 | 125.39 | 6.69 | 0.0103 |
| -2.09 | 0.20 | | 0.06 | | 4 | -58.60 | 125.45 | 6.76 | 0.0100 |
| -2.11 | | | 0.11 | 0.08 | 4 | -58.97 | 126.18 | 7.49 | 0.0069 |
| -2.11 | 0.20 | | 0.05 | 0.08 | 5 | -58.56 | 127.49 | 8.79 | 0.0036 |

**Table S6** Complete negative binomial model set for predicting fluke intensity. Blank values are omitted from the model. Model coefficients are presented under the variable. All models predict fluke intensity ~ present variables

| (Intercept) | Low vegetation | Liver mass | PC1 | PC2 | df | logLik | AICc | delta | weight |
|---|---|---|---|---|---|---|---|---|---|
| 0.27 | | 1.08 | 0.43 | | 5 | -278.51 | 567.39 | 0.00 | 0.4839 |
| 0.27 | 0.07 | 1.07 | 0.41 | | 6 | -278.31 | 569.15 | 1.76 | 0.2006 |
| 0.27 | | 1.07 | 0.44 | 0.08 | 6 | -278.31 | 569.15 | 1.76 | 0.2003 |
| 0.27 | 0.07 | 1.06 | 0.42 | 0.09 | 7 | -278.09 | 570.88 | 3.50 | 0.0843 |
| 00.28 | | 1.06 | | | 4 | -283.20 | 574.64 | 7.26 | 0.0129 |
| 0.27 | 0.17 | 1.05 | | | 5 | -282.39 | 575.16 | 7.77 | 0.0099 |
| 0.28 | | 1.06 | | 0.03 | 5 | -283.18 | 576.73 | 9.34 | 0.0045 |
| 0.26 | 0.17 | 1.04 | | 0.04 | 6 | -282.35 | 577.23 | 9.84 | 0.0035 |
| 0.63 | | | 0.35 | 0.30 | 5 | -307.85 | 626.07 | 58.68 | 8.77E-14 |
| 0.62 | 0.14 | | 0.30 | 0.30 | 6 | -307.25 | 627.02 | 59.63 | 5.45E-14 |
| 0.67 | | | 0.32 | | 4 | -309.92 | 628.10 | 60.71 | 3.18E-14 |
| 0.61 | 0.22 | | | 0.29 | 5 | -309.45 | 629.27 | 61.88 | 1.77E-14 |
| 0.66 | 0.12 | | 0.28 | | 5 | -309.47 | 629.32 | 61.93 | 1.72E-14 |
| 0.63 | | | | 0.28 | 4 | -310.78 | 629.81 | 62.42 | 1.35E-14 |
| 0.66 | 0.19 | | | | 4 | -311.28 | 630.81 | 63.42 | 8.19E-15 |
| 0.67 | | | | | 3 | -312.39 | 630.94 | 63.55 | 7.69E-15 |

**Table S7** Complete negative binomial model set for predicting bladderworm intensity. Blank values are omitted from the model. Model coefficients are presented under the variable. All models predict bladderworm intensity ~ present variables

| (Intercept) | Low vegetation | Liver mass | PC1 | PC2 | df | logLik | AICc | delta | weight |
|---|---|---|---|---|---|---|---|---|---|
| -1.09 | | 0.64 | | | 4.00 | -121.58 | 251.42 | 0.00 | 0.2550 |
| -1.12 | | 0.75 | | -0.31 | 5.00 | -120.87 | 252.10 | 0.69 | 0.1807 |
| -1.11 | | 0.72 | 0.23 | | 5.00 | -121.05 | 252.48 | 1.07 | 0.1496 |
| -1.10 | 0.02 | 0.65 | | | 5.00 | -121.58 | 253.53 | 2.11 | 0.0887 |
| -1.14 | | 0.79 | 0.18 | -0.26 | 6.00 | -120.56 | 253.65 | 2.23 | 0.0835 |
| -1.12 | -0.01 | 0.75 | | -0.31 | 6.00 | -120.86 | 254.25 | 2.84 | 0.0617 |
| -1.11 | -0.06 | 0.72 | 0.25 | | 6.00 | -121.02 | 254.57 | 3.16 | 0.0526 |
| -0.94 | | | | | 3.00 | -124.52 | 255.18 | 3.77 | 0.0388 |
| -1.14 | -0.07 | 0.80 | 0.20 | -0.27 | 7.00 | -120.51 | 255.73 | 4.32 | 0.0294 |
| -0.95 | | | 0.08 | | 4.00 | -124.44 | 257.13 | 5.71 | 0.0146 |
| -0.95 | | | | -0.09 | 4.00 | -124.45 | 257.14 | 5.72 | 0.0146 |
| -0.94 | 0.01 | | | | 4.00 | -124.52 | 257.28 | 5.87 | 0.0136 |
| -0.95 | | | 0.06 | -0.07 | 5.00 | -124.40 | 259.18 | 7.76 | 0.0053 |
| -0.95 | -0.02 | | 0.09 | | 5.00 | -124.44 | 259.25 | 7.83 | 0.0051 |
| -0.95 | 0.00 | | | -0.09 | 5.00 | -124.45 | 259.26 | 7.85 | 0.0050 |
| -0.95 | -0.02 | | 0.07 | -0.07 | 6.00 | -124.40 | 261.32 | 9.90 | 0.0018 |
